# SilentMutations (SIM): a tool for analyzing long-range RNA-RNA interactions in viral genomes and structured RNAs

**DOI:** 10.1101/424002

**Authors:** Daniel Desirò, Martin Hölzer, Bashar Ibrahim, Manja Marz

**Affiliations:** European Virus Bioinformatics Center, Jena, Germany; RNA Bioinformatics and High-Throughput Analysis Jena, Friedrich Schiller University Jena, Jena, Germany; Leibniz Institute for Age Research-Fritz Lipmann Institute, Jena, Germany

**Keywords:** Codon Mutation, Virology, RNA virus, Virus Bioinformatics, RNA secondary structure, double-mutant, silent mutation

## Abstract

**Background:** A single nucleotide change in the coding region can alter the amino acid sequence of a protein. In consequence, natural or artificial sequence changes in viral RNAs may have various effects not only on protein stability, function and structure but also on viral replication.

In recent decades, several tools have been developed to predict the effect of mutations in structured RNAs such as viral genomes or non-coding RNAs. Some tools use multiple point mutations and also take coding regions into account. However, none of these tools was designed to specifically simulate the effect of mutations on viral long-range interactions.

**Results:** Here, we developed `SilentMutations (SIM)`, an easy-to-use tool to analyze the effect of multiple point mutations on the secondary structures of two interacting viral RNAs. The tool can simulate disruptive and compensatory mutants of two interacting single-stranded RNAs. This allows a fast and accurate assessment of key regions potentially involved in functional long-range RNA-RNA interactions and will eventually help virologists and RNA-experts to design appropriate experiments.

`SIM` only requires two interacting single-stranded RNA regions as input. The output is a plain text file containing the most promising mutants and a graphical representation of all interactions.

**Conclusion:** We applied our tool on two experimentally validated influenza A virus and hepatitis C virus interactions and we were able to predict potential double mutants for *in vitro* validation experiments.

**Availability:** The source code and documentation of `SIM` are freely available at github.com/desiro/silentMutations.

## Introduction

In the last decades, several computational tools for the analysis of RNA secondary structures have been developed. However, tools specifically targeting virus needs are still rare and under-researched^1–3^. For example, long-range RNA-RNA interactions (LRIs) play an important role in the life cycle of RNA viruses.

Many LRIs are already known to act directly as activators or inhibitors of viral replication and translation^4^. For example, several interactions have been recently identified as potential new LRIs in the genome of the hepatitis C virus (HCV) using `LRIscan`^5^. Some of these computationally identified interactions have already been verified experimentally^6–13^.

Apart from long-range interactions forming in a single sequence (intra-molecular structures), LRI-like structures can also occur between two separate sequences, for example between two viral segments. Whereas the general function of such interactions is still under investigation, they can be seen as viral LRIs and are presumably responsible for the correct packaging of all segments into the viral capsid of segmented viruses such as the influenza A virus (IAV)^14–17^.

Since the number of known and possibly functional LRIs is growing rapidly, it is essential to have an effective verification method. Technically, such interactions can be destroyed by removal or mutation of the interacting segments using *in vitro* experiments^8,18^. Such sequence changes can result in secondary structure changes and finally manifest different viral titers. However, with the alteration of the sequence, unwanted effects can arise that not only effect the LRI.

To cope with this issue, we can modify both interacting RNA segments simultaneously. Additionally, we aim for the combination of both mutated segments to result in a similar interaction strength in comparison to the interaction between the wild type (WT) sequences. A single mutated sequence segment in combination with the respective WT sequence should destroy the interaction. Such a technique was recently used to verify a possible LRI between two IAV segments^15^.

Several computational tools are available that can alter a sequence and report alternative secondary structures with one or even multiple mutations^19–26^. However, these tools are designed for specific applications and do not meet the requirements of the experimental technique described above: the combinatorial *in vitro* verification of RNA-RNA interactions.

In this study, we present a tool called `SilentMutations (SIM)` that effectively simulates synonymous (silent) compensatory mutations in two single-stranded viral RNAs and is therefore appropriate for the *in vitro* assessment of predicted LRIs.

## Materials and Methods

Here, we present a command-line tool, called `SilentMutations (SIM)`, that can simulate synonymous structure-disrupting and structure-preserving mutation pairs within coding regions for long-range RNA-RNA interaction experiments. The tool has been written in `Python` (v3.6.5) and heavily relies on the `RNAcofold` python site-package of the `ViennaRNA Package` (v2.4)^27^. As the `ViennaRNA Package` is available for Linux, Windows, and MacOS, all three different platforms are supported by `SIM`. The various parameters of `SIM` are fully adjustable and several filters (**Fig. 1**) allow a fast and accurate prediction, therefore the tool can be run on a standard notebook.

**Figure 1:**
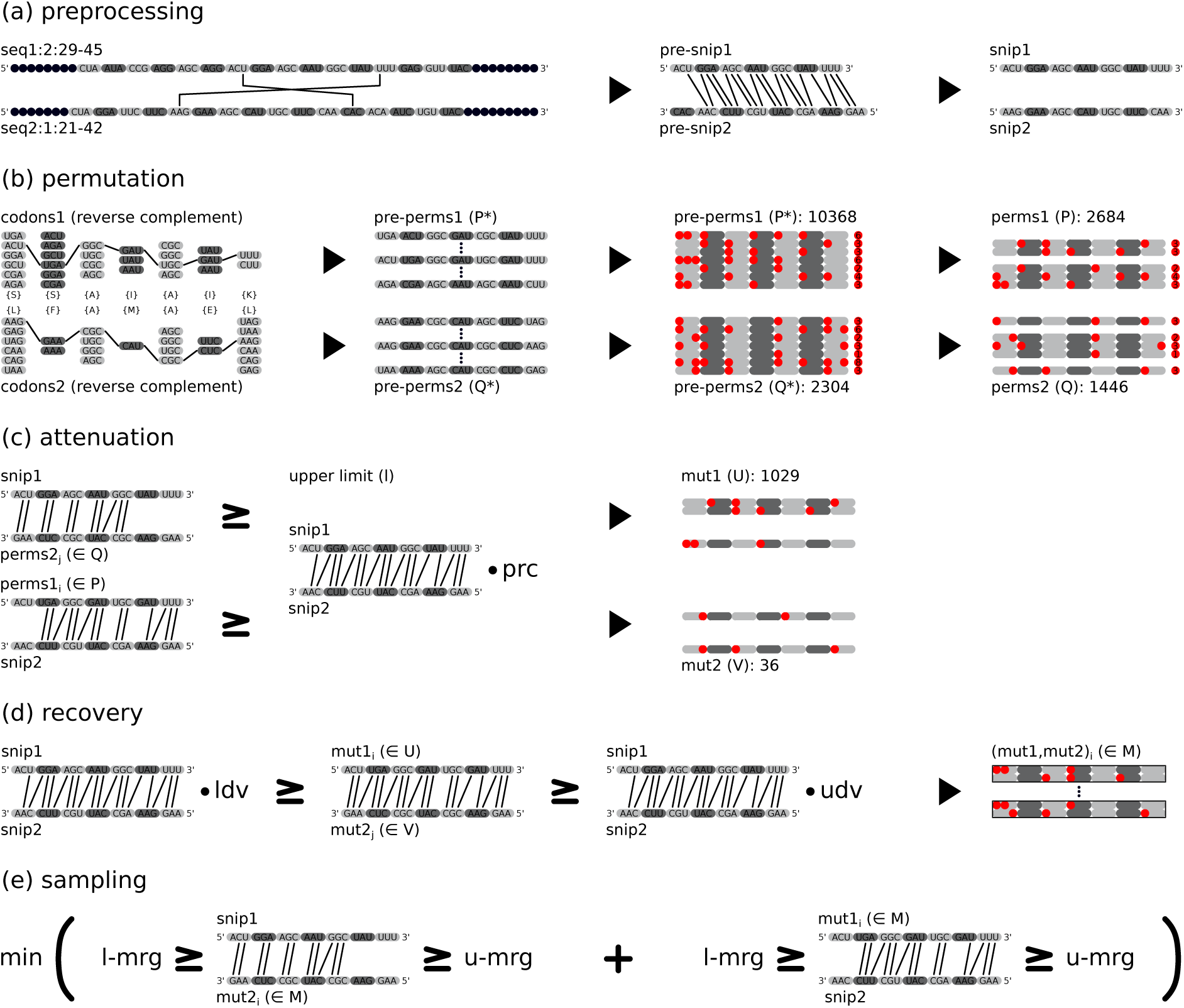
Overall workflow of SIM, exemplarily shown for two sequences from a negative single-stranded RNA virus genome (ssRNA-) **(a)** The first step extracts the defined range in each sequence and possibly increases the range to preserve codons based on the given reading frame. Both so-called *pre-snips* are folded with `RNAcofold` to remove unpaired codons at the endings. We refer to an extracted single-stranded RNA snippet as *snip*. **(b)** All possible codon permutations (here, reverse complements due to ssRNA-) are generated for both *snips* (called *perms*) to conserve the amino acid sequence. Permutations with too many mutations defined by the mutations (`-mut`) parameter are discarded. **(c)** To further decrease the total number of combinations, `SIM` will fold each *snip* with all permutations of the other *snip* and keep permutations with a *mfe* higher or equal to *mfe*(*snip1,snip2*) times the filter percentage (`-prc`) parameter, denoted as upper limit *l*. **(d)** All remaining *snip1* permutations will then be folded against all *snip2* permutations to find *snip1* and *snip2* mutations with a similar fold *mfe*. A double-mutant *mfe*(*mut1,mut2*) fold is considered to be similar, if its *mfe* is lower than the wild-type *mfe*(*snip1,snip2*) times the lower deviation (`-ldv`) parameter and higher than *mfe*(*snip1,snip2*) times the upper deviation (`-udv`) parameter. **(e)** The last step will minimize a combined single-mutant *mfe*(*snip1,mut2*) and *mfe*(*mut1,snip2*) by keeping both mutations in a similar range depending on the mutation range (`-mrg`) parameter. The range for each mutation combination is defined by (*mfe*(*snip1, mut2*) + *mfe*(*mut1, snip2*)) · 0.5 · (1 + `-mrg`) for the lower threshold (l-mrg) and (*mfe*(*snip1, mut2*) + *mfe*(*mut1, snip2*)) · 0.5 · (1 - `-mrg`) for the upper threshold (u-mrg). Different sets and their abbreviations are given in brackets. Details can be found in the Methods.

A simple run takes about 10 seconds when using the standard parameters, two sequences of length 21 as input, and four cores. However, increasing the length of the two sequences to 30nt already results in a runtime of 1 minute and 10 seconds. Generally, the runtime highly depends on the parameter setup and the length of the input sequences.

For two nucleotide sequences of length *k* and *l* and the number of possible synonymous codons *c*, the worst case runtime can be described by 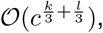 with *c* between 1 and 6 (average: 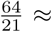 3.048). This large search space can be drastically reduced by various filter steps implemented in `SIM` (see below).

Because the runtime depends exponentially on the length of the input sequences, we recommend to limit the interaction length to maximal 30 nt. Additionally, longer sequences are more likely to form intra-molecular secondary structures instead of forming stable structures with each other. Whereas `SIM` can be used to evaluate longer sequences, the tool is originally intended for two smaller sequence segments forming a stable secondary structure. For the prediction of LRIs in viral RNA genomes and longer sequences, we recommend `LRIscan`^5^. The main computational steps of `SIM` are (1) preprocessing, (2) permutation, (3) attenuation, (4) recovery, and (5) sampling.

A particular challenge is posed by negative single-stranded RNA viruses such as Ebola, Marburg, and Coronaviruses, as most databases only contain the positive strand of the virus genome. While the protein-coding sequence is encoded on the positive strand, the folding of these viral RNAs happens on the negative strand. Depending on the user input, `SIM` can automatically create the reverse-complement strand of negative single-stranded RNA viruses for folding, while maintaining codon integrity on the positive strand. This can be easily achieved with the `-virusclass=ssRNA-`, `-reverse`, and `-complement` options. The user can directly provide the positive strand sequence from the database in a fasta file, with the reading frame and regions on the positive strand.

The default output of `SIM` prints all relevant information of the folding in a plain text file and additionally creates for each folding a `VARNA`^28^ command for visualization. Additionally, if the tool can find installations of `VARNA` and `inkscape`^29^, it will directly generate high-quality vector graphics of the various foldings. A default binary of `VARNA` (v3.93) is provided with the tools source code.

`SIM` is primarily designed for two sequence snippets (hereinafter referred to as snips) in coding regions, but can also be used to simulate interactions between coding and non-coding regions and for interactions between two non-coding regions. This can be done with the `-noncoding1` and `-noncoding2` parameters, to define the first or second sequence as non-coding, respectively. This makes the prediction of interactions between RNA molecules with selection pressure on the sequence for translation and RNA molecules with selection pressure on secondary structures possible.

We use the following notation for the minimum free energy (*mfe*) obtained by folding sequence *x* with sequence *y* via `RNAcofold`:

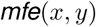

The *mfe* is represented by a negative value in 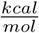 and therefore having a lower *mfe* results in a more stable structure.

### Preprocessing

In the first step of `SIM`, sequence snips are extracted from longer sequence templates while keeping codons intact (**Fig. 1**). Especially, the difficult extraction of snips in the exact reading frame from negative sense single-stranded RNAs can be easily handled by the build-in functionalities of the tool. Thus, `SIM` directly accepts the whole sequence as input, together with a start and end position for the snips and the reading frame. The tool first extracts the requested interacting segments from each sequence. Each of them will be automatically increased in both directions if the provided range would otherwise split a codon, based on the predefined reading frame (**Fig. 1**). This ensures a preservation of all codons. Both sequences, *pre-snip1* (*ps*_1_) and *pre-snip2* (*ps*_2_), will then be fold together using `RNAcofold` to acquire only codons involved in the folding. Unpaired codons at the termini will be removed to get the final *snip1* (*s*_1_) and *snip2* (*s*_2_) from which `SIM` calculates the two mutant sequences *mut1* (*m*_1_) and *mut2* (*m*_2_).

### Permutation

In this step, all possible permutations *pre-perms1* (*P*^*^) and *pre-perms2* (*Q*^*^) of synonymous codons are created from each sequence in order to find the most suitable mutations that maximize the distance between *mfe*(*s*_1_,*s*_2_) and *mfe*(*s*_1_,*m*_2_) and the distance between *mfe*(*s*_1_,*s*_2_) and *mfe*(*m*_1_,*s*_2_), while keeping the wild-type *mfe*(*s*_1_,*s*_2_) and the double-mutant *mfe*(*m*_1_,*m*_2_) similar. The number of these permutations can be vast, depending on the number of synonymous codons and the length of the sequence. To reduce the computational complexity, this step also removes every permutation sequence with more mutations than predefined by the `-mutations` parameter resulting in the sets *perms1* (*P*) and *perms2* (*Q*), see **Fig. 1**. With our implementation of this parameter, we aim to keep the number of introduced mutations small, while simultaneously maximizing the *mfe* difference between wild-type and single mutants.

### Attenuation

In this step, the computational complexity is further reduced and valid sequences with reduced *mfe* folding scores are created. Importantly, having a small *mfe* difference between the wild-type (*mfe*(*s*_1_,*s*_2_)) and the single mutants (*mfe*(*s*_1_,*m*_2_) or *mfe*(*m*_1_,*s*_2_)) would diminish the mutational effect on the interaction. It is therefore vital to increase the distance between these *mfe* scores. Therefore, the user can directly specify the required distance with the filter percentage (`-filterperc`) parameter. The defined value is then multiplied with the *mfe*(*s*_1_,*s*_2_) value to obtain an upper limit (denoted *l*) for *mfe*(*s*_1_,*m*_2_) and *mfe*(*m*_1_,*s*_2_). Then, the tool creates all foldings between *s*_1_ and *Q* as well as *P* and *s*_2_. Each folding *mfe*(*s*_1_, *q_j_* ∈ *Q*) and *mfe*(*p_i_* ∈ *P,s*_2_) (with *i* and *j* defining the i-th and j-th element of set *P* and *Q*, respectively) is only considered for the next recovery step if the *mfe* is higher than *l*. The new sets *mut1* (*U*) and *mut2* (*V*) have a drastically reduced size compared to *P* and *Q* which greatly reduces the computational complexity for finding the two mutant sequences *m*_1_ and *m*_2_ where *mfe*(*m*_1_,*m*_2_) is similar to the wild-type *mfe*(*s*_1_,*s*_2_). While it seems to be convenient to set a very low `-filterperc` parameter, this could also result in an empty permutation set. On the other hand, setting a high `-filterperc` could result in longer running times. It is therefore recommended to start with the default *0.5* and later adjust this parameter depending on the performance.

### Recovery

To find double-mutants (mut) with a similar folding *mfe* such as the wild-type (WT) folding *mfe*, each sequence *u_i_* ∈ *U* and *v_i_* ∈ *V* has to be folded against each other with `RNAcofold`. The running time of this folding process strongly depends on the previous attenuation step. When using the unprocessed *P^*^* and *Q^*^* sets from the example in **Fig. 1**, the number of required folding operations would be around 24 million and reduced to only 37,000 for the filtered sets *U* and *V*. Some LRIs function as inhibitors for viral replication^4^ and a stronger interaction would decrease functional viral reproduction. For the simulation of the double-mutant interaction it is therefore necessary to set a lower but also an upper similarity threshold. Each of the calculated *mfe*(*u_i_* * *U,v_j_* ∈ *V*) values are therefore compared with the wild-type *mfe*(*s*_1_,*s*_2_). A combination of two sequences *u* and *v* is only valid if

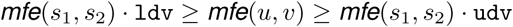

with `ldv` as the lower deviation (`-lowerdev`) and `udv` as the upper deviation (`-upperdev`) parameter. Valid (*u,v*) double-mutant pairs are stored as tuples in set *M*.

### Sampling

Now, all pairs in the final set *M* contain only mutated sequence pairs with the desired properties. However, greedily selecting the i-th tuple (*u*, *v*)_*i*_ ∈ *M* that minimizes

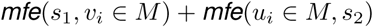

could result in unfavorable interaction strengths between WTs and mutants. An unfavorable *v* and *u* pair would result in a significantly higher single-mutant *mfe* of the mutant folded with the WT sequence in comparison to the *mfe* of the other mutant and WT combination. Therefore, it is recommended to keep both, *mfe*(*s*_1_,*v_i_* ∈ *M*) and *mfe*(*u_i_* ∈ *M,s*_2_), in a similar range while minimizing the combined *mfe*. Accordingly, the sampling step takes the mutation range (`-mrg`) parameter into account and calculates the lower threshold `l-mrg` as

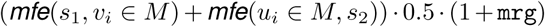

and the upper threshold `u-mrg` as

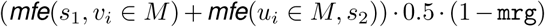

for each mutant tuple. The most suitable mutant pair, holding all predefined requirements, is finally found by calculating:

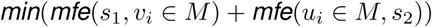

with

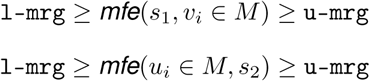

## Results and Discussion

`SIM` was primarily developed as a tool for designing synonymous mutations required for probing LRIs detected in viral genomes, such as previously performed by Gavazzi *et al.*in influenza A viruses *in vitro*^15^. With the help of `SIM`, experimentalists can simulate for the first time specific RNA-RNA interactions of viral coding-and non-coding sequence snippets, potentially forming stable long-range interactions. Using `SIM`, the search space of possible double-mutant sequences, forming a stable secondary structure with a comparable *mfe* to the wild-type structure and simultaneously disrupting the single-mutants structure, can be drastically reduced. Therefore, `SIM` can be used prior time - and cost-consuming wet-lab experiments to simulate promising double-mutant sequences, that preserve the wild-type LRI but are not functional as single-mutants. In this context, the use of `SIM` is not limited to RNA interactions between viral sequences, as the tool can be used for calculations between any two interacting RNA sequences. To validate *SIM* and to show that the tool is able to predict biological relevant mutants, we used two experimentally verified interactions, one in the influenza A virus H5N2 A/finch/England/2051/1991 strain and another in the hepatitis C virus type 1b strain (accession: AJ238799.1).

### Application to influenza A virus

The influenza A virus (IAV) genome consists of 8 viral ribonucleoproteins (vRNPs) and each vRNP segment includes one of 8 different negative sense and single-stranded viral RNAs. It is hypothesized, that these segments are packed selectively through RNA-RNA interactions between the 8 segments^16,30–32^. To test this, Gavazzi *et al.*^15^ conducted an *in vitro* mutation experiment in IAV, which is perfectly suited to validate our tool. They took a yet unconfirmed interaction from one of their previous experiments^14^ between the PB1 and the NS segment and introduced four trans-complementary point substitutions by hand. An interaction between the PB1 mutant and NS mutant resulted in a similar viral titer than using the PB1 WT and NS WT segment. The introduction of only the PB1 mutant or NS mutant led in both cases to an attenuation of the viral replication.

To show that we are able to obtain the same experimental results as Gavazzi *et al*.^15^ computationally, we adjusted some key parameters of `SIM`. Importantly, we limited the number of possible mutation pairs to four and calculated the *mfe* between the WT and mutant (mut) combinations in mere seconds. We further increased the search space of single mutants through setting the filter percentage parameter to 0.6, derived from the difference between the lowest single-mutant and the WT *mfe* value 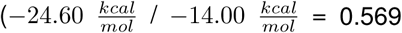; see **Fig. 2 a and c**). Because there exist multiple double mutants with reasonable *mfe* values, we also had to define an exact search range for the double-mutant. The lower deviation was set to 1.07 and the upper deviation to 1.08, based on the difference between the double-mutant and WT *mfe* values 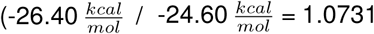; see **Fig. 2 a and b**). The `SIM` results of the secondary structure predictions and *mfe* values for the two IVA segment combinations are shown in **Fig. 2**. By only allowing a maximum of four mutation pairs, we were able to calculate the same structures and mutations as previously proposed by Gavazzi *et al*.^15^. Moreover, our results show a *mfe* difference of 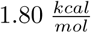 between wild-type and double-mutant (**Fig. 2 a and b**) and a *mfe* difference of 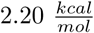 between single-mutants (**Fig. 2 c and d**). Again, we want to point out that for the *in silico* experiment it is preferable to have a similar *mfe* between the two wild-type IAV segments *mfe*(*NS_WT_*, *PB1_WT_*) and the double-mutant *mfe*(*NS_mut_*, *PB1_mut_*) as well as between the single mutants *mfe*(*NS_WT_*, *PB1_mut_*) and *mfe*(*NS_mut_*, *PB1_WT_*).

**Figure 2:**
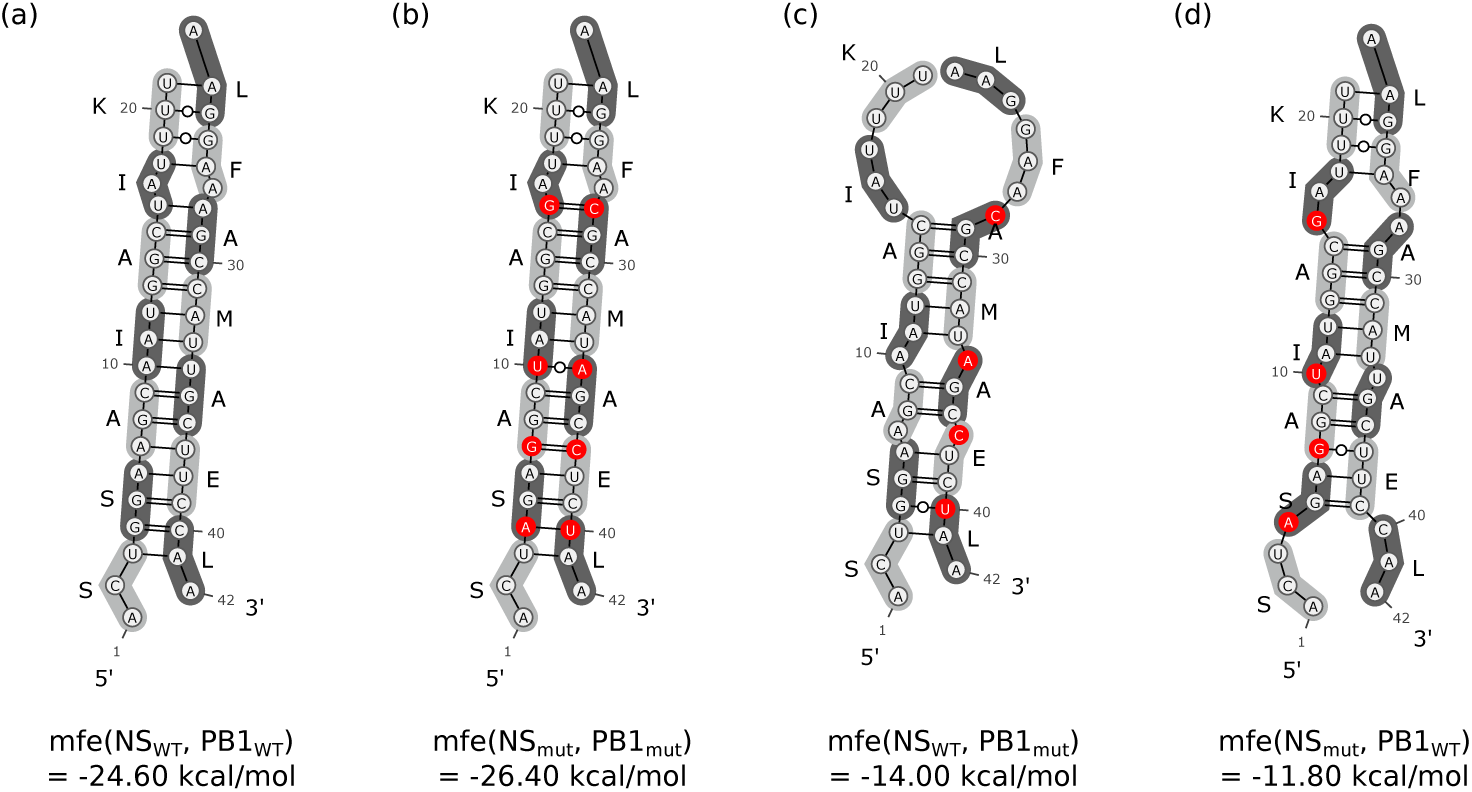
IAV interactions like previously shown by Gavazzi *et al*. We adjusted the parameters of `SIM` and allowed only four mutation pairs in maximum to calculate the *mfe* values for mutated sequence combinations of the NS and PB1 segments like previously shown by Gavazzi *et al*.^15^. The *mfe* values and secondary structures are shown for **(a)** the wild-type interaction, **(b)** the double-mutant interaction, **(c)** the first single-mutant (PB1_*mut*_) interaction, and **(d)** the second single-mutant (NS_*mut*_) interaction. This visualization was automatically generated by `SIM` using `VARNA`.

In a next step and by using `SIM` with default parameters, we were able to calculate a double-mutant that not only reflects the results of Gavazzi *et al*.^15^, but also shows slightly lower *mfe* differences between the wild-type and the double-mutant 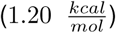 as well as between the two single-mutants 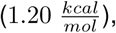 see **Fig. 3**.

**Figure 3:**
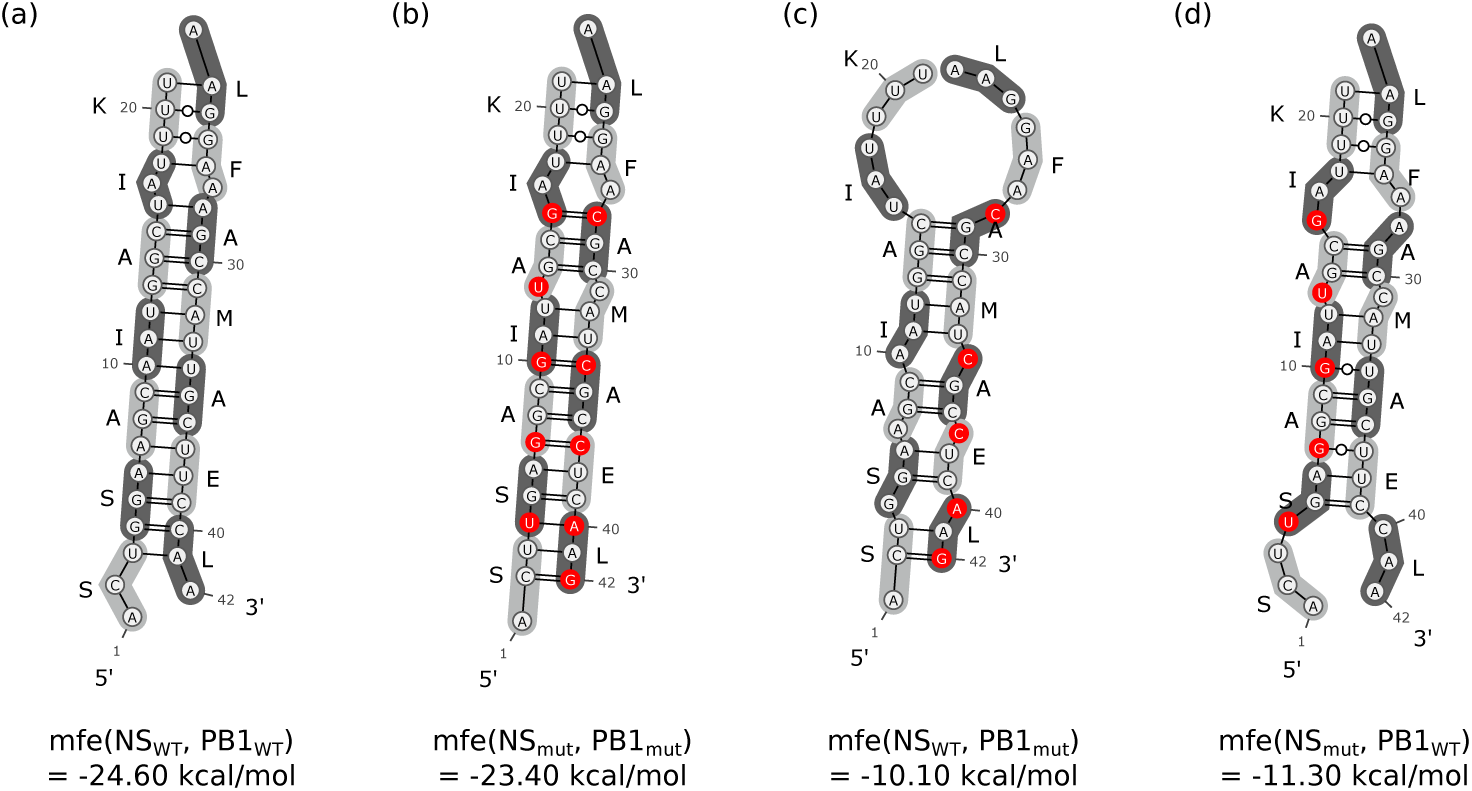
IAV interactions calculated by SIM. By applying `SIM` to the same IAV sequence snips of the NS and PB1 segments, we were able to show lower *mfe* differences between **(a)** the wild-type interaction and **(b)** the double-mutant interaction, as well as between the two single-mutant interactions **(c,d)**.

Therefore, the interaction strengths of *mfe*(*NS_WT_*, *PB1_WT_*) and *mfe*(*NS_mut_*, *PB1_mut_*), as well as *mfe*(*NS_WT_*, *PB1_mut_*) and *mfe*(*NS_mut_*, *PB1_WT_*) are more closely to each other in comparison to the results of Gavazzi *et al*.^15^. As a possible disadvantage, our simulation had to introduce another point mutation into each single-stranded RNA snip (**Fig. 3**).

### Application to hepatitis C virus

The hepatitis C virus (HCV) genome consists of a positive single-stranded RNA of about 10kb in length. This RNA is translated into a single polyprotein that is later cleaved into four structural (C, E1, E2, p7) and six nonstructural (NS2, NS3, NS4A, NS4B, NS5A, RdRp) proteins by viral and host proteases^33,34^. Both UTR regions of the viral genome are highly structured and have been extensively studied in the past^33,35–38^.

To validate our tool, we have chosen a well-studied interaction^8^ between the 3’UTR and the end region of the ORF encoding the HCV polyprotein. Several studies^9,33,39–42^ have already shown that the RNA replication highly depends on the conserved structures of the X-tail^33,43^ sequence contained in the 3’UTR. This sequence is presumed to contain three experimentally verified stem-loops (SLI, SLII, SLIII)^44^ which may interact with other segments of the viral genome through long-range RNA-RNA interactions to regulate replication^40^. The LRI between the free nucleotides of the hairpin from the 3’ SLII structure and the free nucleotides of the hairpin from the 5BSL3.2 structure have been investigated in many HCV studies.

The interaction was first verified by Friebe *et al*.^8^ and was also computationally predicted with `LRIscan` by Fricke *et al*.^5^. Furthermore, we have chosen this interaction as an example, since on interacting segment is located in a coding region and the other segment in a non-coding region. Applying `SIM` on this interaction resulted in two compensatory point mutations in each snip (**Fig. 4**). As the 3’SLII region is non-coding, only the point mutations in the snip from the 5BSL3.2 structure are silent. Our results show, that the wild-type structure *mfe*(*5BSL3.2_WT_*, *SLII_WT_*) should have a similar strength compared to the calculated double-mutant structure *mfe*(*5BSL3.2_mut_*, *SLII_mut_*), see **Fig. 4**. Additionally, both *mfe*(*5BSL3.2_WT_*, *SLII_mut_*) and *mfe*(*5BSL3.2_mut_*, *SLII_WT_*) weaken the interaction significantly.

**Figure 4:**
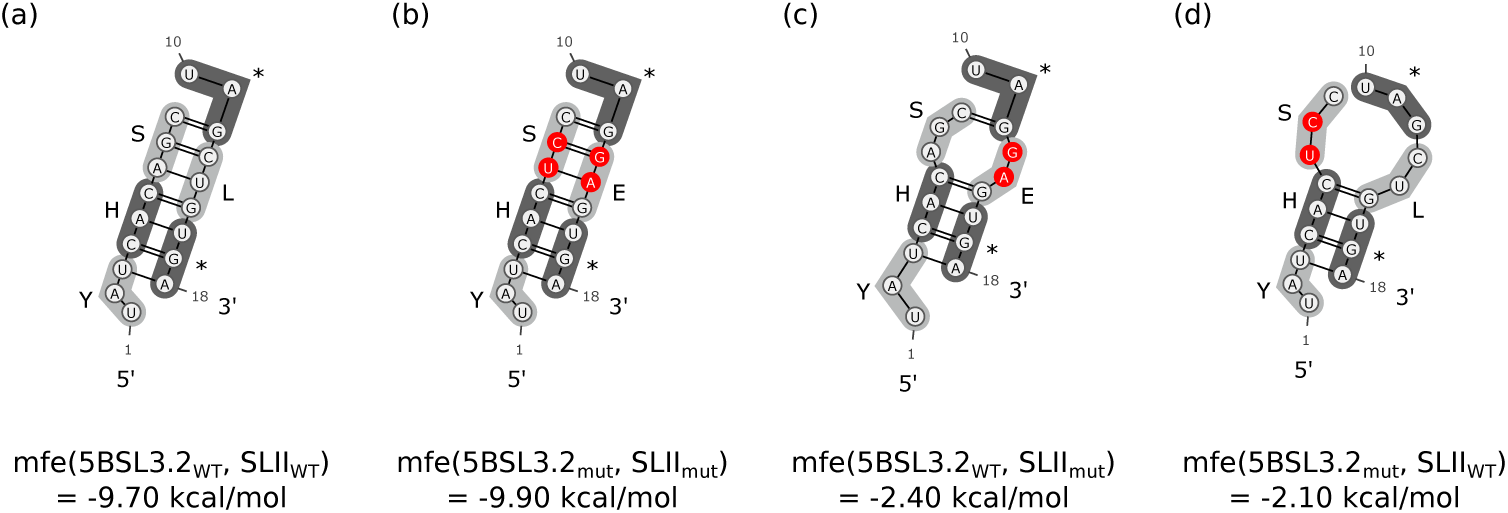
HCV interaction calculated with SIM. We used `SIM` to calculate a double-mutant interaction between a HCV protein-coding snip of 5BSL3.2 and an long-range interacting non-coding snip of the 3’ SLII region. Again, our calculations show similar *mfe* values between the stable wild-type and double-mutant structures **(a,b)** as well as between the structure-destroying and less stable single-mutants **(c,d)**.

We suggest, that our simulated mutations in HCV may be used to verify the given long-range interaction. Taking all the new long-range interactions found by Fricke *et al*.^5^ into account, we propose that our tool can be used to create mutation experiments for every predicted LRI to provide evidence for a biological function of these interactions. Such experiments would be especially interesting for IAV, where the exact packaging process of the vRNP segments is not yet fully understood.

### Conclusion

With the development of `SIM`, we offer a simple and fast way to analyze possible interactions between vRNPs and other RNA molecules. The tool can be used to heavily reduce the search space of possible synonymous mutation interactions between two RNAs. Another difficulty when creating silent IAV mutations lies in preserving the codons on the positive strand, while mutating the negative strand, and is also intercepted by `SIM`. Furthermore, our tool provides a significant speedup, not only in the verification of interactions in these two viruses, but also for many other virus families. Our simulations will help to gather a deeper understanding of the translation and replication processes in viruses and also how long-range interactions are regulating these.

A promising approach for the future would be the combined use of `LRIscan` and `SIM` to detect previously unknown LRIs and at the same time perform possible mutational verification experiments for each LRI.

## Author Contributions

DD performed the design, development and programming of the tool and wrote the main draft of the paper. MH, BI, and MM contributed in writing, discussions and proofreading of the final manuscript. All authors read and approved the final manuscript.

